# Side-by-side evaluation of two mouse models for Crimean-Congo Hemorrhagic Fever Virus infection

**DOI:** 10.1101/2025.08.20.671421

**Authors:** Cornelius Rohde, Anke-Dorothee Werner, Michelle Gellhorn Serra, Petra Emmerich, Nataša Knap, Markus Eickmann, Stephan Becker, Verena Krähling, Alexandra Kupke, BSL4 Animal Facility Team

## Abstract

Crimean-Congo hemorrhagic fever virus (CCHFV) is the causative agent of a severe hemorrhagic fever in humans, associated with case fatality rates ranging from 10 to 40%. Due to the lack of approved vaccines or specific antiviral treatments, CCHFV is classified as a biosafety level 4 (BSL4) pathogen in most countries and designated a priority pathogen by the World Health Organization (WHO). To facilitate the preclinical assessment of medical countermeasures, we have established two murine models using C57BL/6J IFNAR^−/−^ mice, which lack the IFNα/β receptor, infected with the phylogenetically distinct CCHFV strains Afghanistan09-2990 (Afg09) and Kosovo Hoti (Hoti). Infection with both CCHFV strains in IFNAR^−/−^ mice resulted in significant weight loss, with Afg09 infection leading to more severe clinical disease. Quantitative analysis of viral RNA revealed widespread viral dissemination across multiple organs in both models. Detection of infectious virus varied by organ and strain. These results confirm and extend previous findings, providing a deeper understanding of CCHFV strain-specific pathogenesis in IFNAR^−/−^ mice. Thereby, these mouse models represent valuable tools for the evaluation of antiviral therapeutics and vaccine candidates, enabling the investigation of cross-lineage protection against genetically diverse CCHFV isolates.

## Introduction

CCHFV is mainly transmitted by tick bites to humans and can cause severe hemorrhagic fever, with 1,000-2,000 fatalities per year worldwide(1). Due to the widespread distribution of the main vector (ticks of the genus *Hyalomma*), CCHFV transmission has been reported in Europe, Africa and Asia. This broad geographic spread is reflected in classification of the virus into five distinct genetic lineages (I-V)(2). In the absence of approved antiviral therapeutics or vaccines, CCHFV is listed as a priority pathogen for research by the WHO, capable of causing a Public Health Emergencies of International Concern (PHEIC)(3, 4).

Preclinical animal models are still required to investigate the efficacy of therapeutics and vaccines against emerging viruses such as CCHFV. For this purpose, transgenic rodent models with defects in their type I interferon response are widely used. In contrast to wild type mice, those animals develop a lethal disease after CCHFV infection. IFNAR^-/-^ or STAT1^-/-^ mice, which lack the signal transducer and activator of transcription 1, are suitable transgenic mouse lines for efficacy studies(5, 6). In addition, wild type mice whose interferon response is downregulated with monoclonal antibodies succumb to CCHFV infection(3, 7). To prepare for studies that aim at evaluating cross-protection capacities of future CCHFV vaccines, it was interesting to evaluate the disease development of phylogenetically differently CCHFV strains in the same mouse model.

In the present study we infected C57BL/6J IFNAR^-/-^ mice with CCHFV Afg09 (Lineage IV, Asia) or CCHFV Hoti (Lineage V, Europe) and compared infection characteristics. Differences in replication kinetics and severity of the disease were monitored.

## Results

In this study, we established a mouse model for CCHFV Afg09 and Hoti, respectively, and analyzed them side-by-side to compare the pathogenesis of both CCHFV isolates. For this purpose, C57BL/6J IFNAR^-/-^ mice were infected intraperitoneally (i.p.) with 100 TCID_50_ of the Afg09 or Hoti isolate. Equal numbers of female and male mice were used to take into account possible sex-specific effects on the clinical outcome. Body weight, body temperature as well as spontaneous behaviour and body condition were assessed at least once daily, to determine a clinical score. Mice with a clinical score of ≥10 or if a clinical score of ≥6 was observed on two consecutive days were sacrificed. Mice infected with CCHFV Afg09 reached the clinical endpoint between days 5-8 post infection (p.i.) (Fig. 1A) with an average clinical score of 24.6 because of weight loss (Fig. 1B), altered behaviour and appearance, and in some cases severely reduced body temperature (Fig. 1C). The course of the Afg09-infection in two mice was so fulminant that they were found dead on day 4 p.i.. Mice infected with CCHFV Hoti reached the humane clinical endpoint at days 4-6 p.i. (Fig. 1A) with an average clinical score of 11.2, which again correlated with loss of body weight (Fig. 1B), altered behaviour, and appearance. In both models, an increase in body temperature (average peak increase of body temperature: Afg09 1.18°C, Hoti 1.28°C) was observed 1–2 days prior to reaching the clinical endpoint. At the clinical endpoint, body temperature declined again, dropping well below the physiological range in some Afg09-infected mice (Fig. 1C), serving as an additional indicator of disease progression.

**Figure 1.**
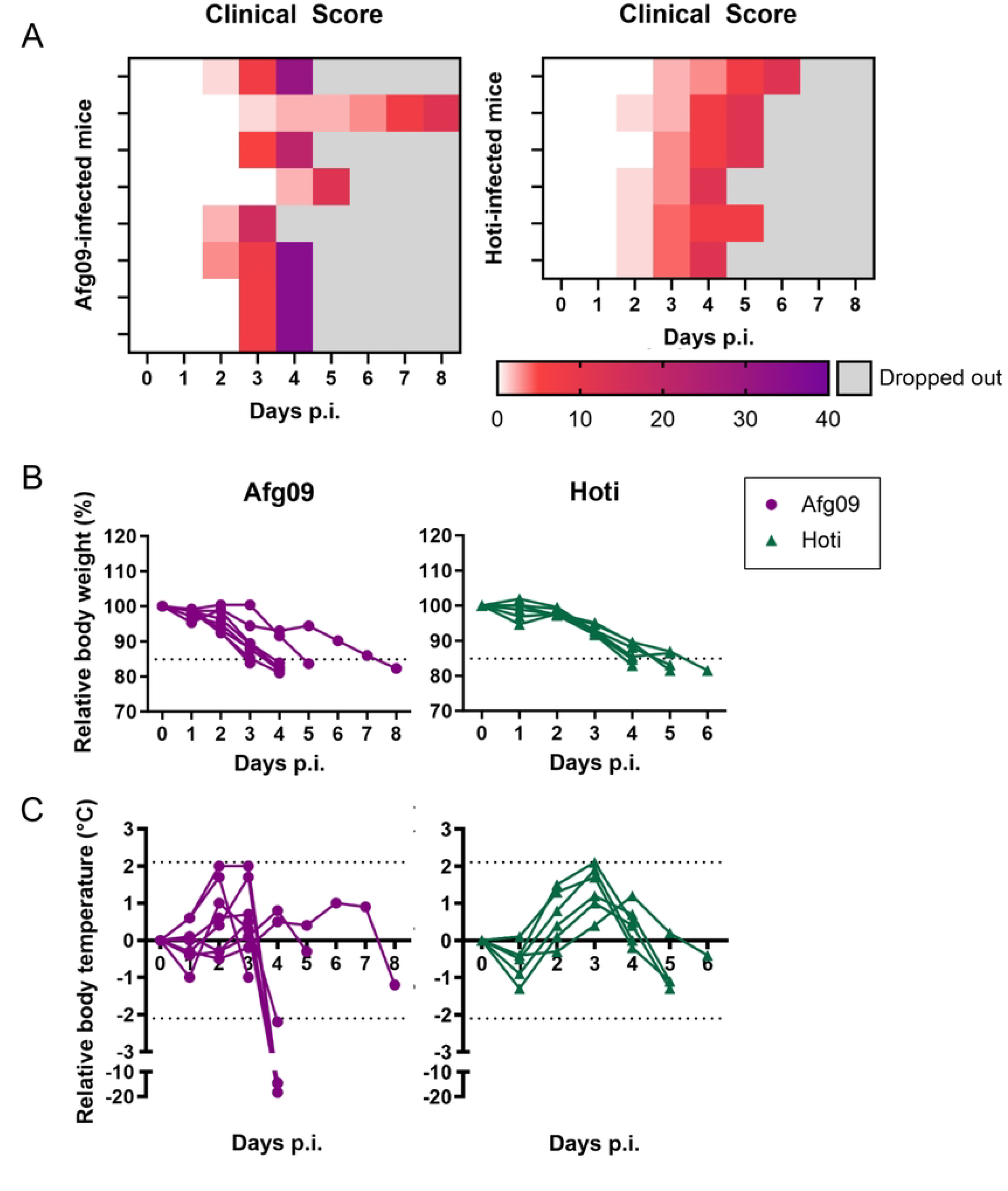
Clinical course of CCHFV Afg09 and Hoti-infected IFNAR^-/-^ mice. IFNAR^-/-^ mice were infected with either 100 TCID_50_ CCHFV Afg09 (purple, n = 8) or Hoti (green, n = 6). The clinical score **(A)**, summarizing appearance, behaviour, body weight **(B)**, and body temperature **(C)**, was monitored daily. After reaching the clinical endpoint (score of ≥10 or ≥6 on two consecutive days), the mice were sacrificed or found dead (grey). Dotted lines mark the clinical endpoints.

During the study, blood was taken from all mice on day 3 p.i. and final blood samples were collected from anaesthetised mice before they were euthanized and organs harvested. To detect CCHFV RNA in organs and serum, qRT-PCR analysis were performed. Virus-specific RNA was detected in serum at day 3 p.i. and at the clinical endpoint, as well as in spleen, liver, central nervous system (CNS), lymph nodes, kidney, eye, thymus, lung, ovaries, testis, and seminal vesicles (Fig. 2A). The amount of infectious CCHFV in organs and final serum samples was determined by TCID_50_ assays. For Afg09, it was shown that infectious virus was detectable in almost all organs and sera until day 5 p.i. for the majority of mice. Viral titers differed among organs, with the highest viral load detected in the liver and the lowest in the seminal vesicles. Low titers of infectious viruses were detected in individual animals in the CNS and the eyes. For Hoti, infectious virus could only be detected in in final sera, thymus, lung and ovaries or testicles (Fig. 2B).

**Figure 2.**
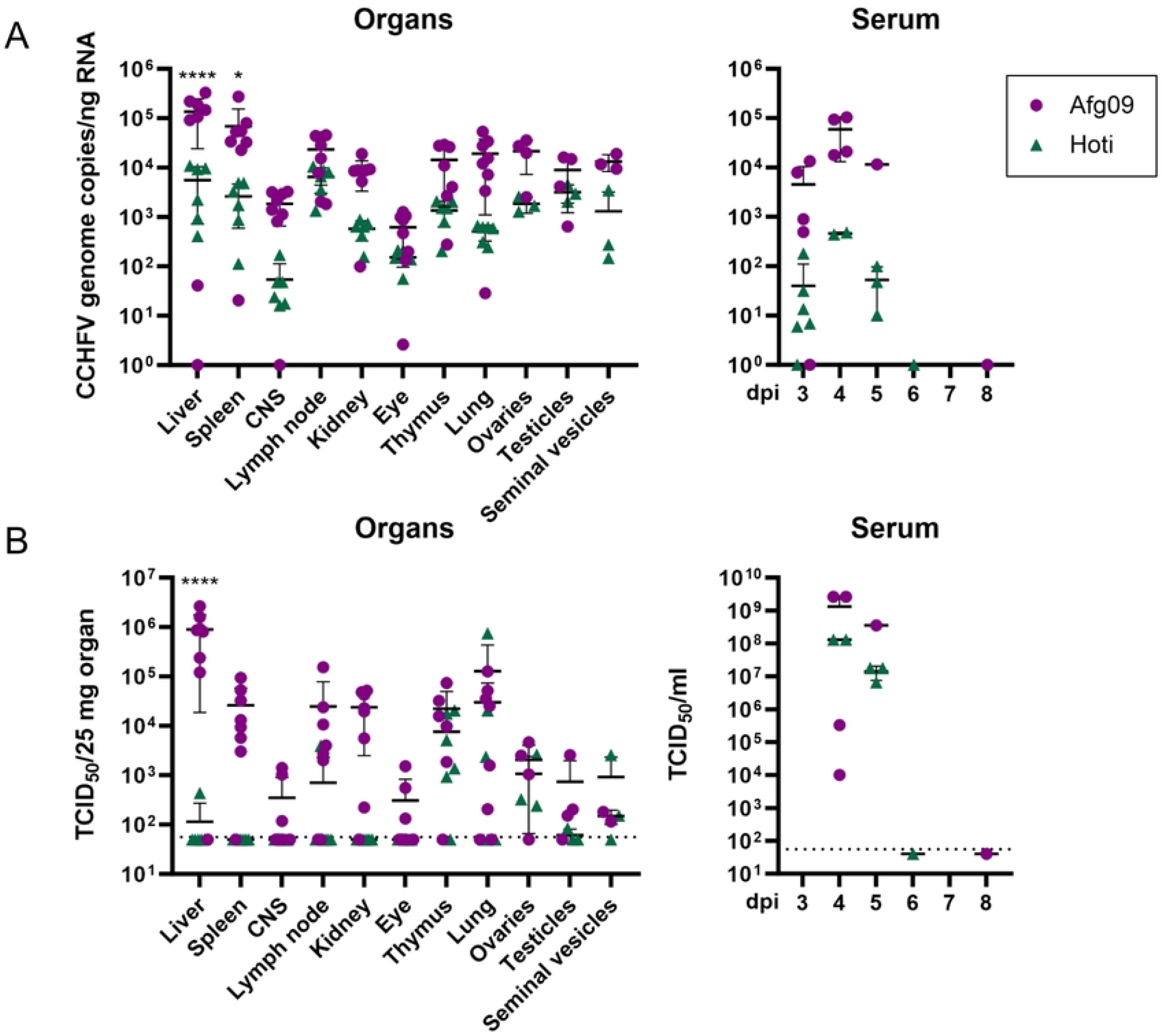
Detection of CCHFV in IFNAR^-/-^ mice infected with Afg09 or Hoti. IFNAR^-/-^ mice were infected with either 100 TCID_50_ CCHFV Afg09 (purple, n = 8) or Hoti (green, n = 6). **A:** CCHFV genome copies were measured by RT-qPCR in organs and serum samples (day 3 and final) of individual mice. **B:** Infectious CCHFV were determined by TCID_50_ assay in organs and final serum samples of individual mice. Each data point represents a sample from an individual animal, data are shown as the means ± SD. Datasets were analyzed using the Šídák’s multiple comparisons test (A) or Tukey’s multiple comparisons test (B). Asterisks indicate statistical significance as detailed between CCHFV Afg09 and Kosovo Hoti group: ∗p ≤ 0.05; ∗∗∗∗p < 0.0001.

The humoral immune response against CCHFV was tested in clinical endpoint sera using a CCHFV whole virus ELISA, detecting mainly NP-specific antibodies, and a CCHFV glycoprotein Gc-specific ELISA. Sera from six infected animals per group were available for analysis. Gc-specific antibodies were first detected at day 4 p.i. in a mouse infected with CCHFV Afg09. Among the four animals that reached the clinical endpoint at day 5 p.i., three mice infected with the Hoti exhibited Gc-specific antibodies in their sera, whereas the Afg09-infected animal did not. Of the two animals that survived beyond day 5, one infected with Hoti (surviving until day 6) and one with Afg09 (surviving until day 8), both developed a broader humoral immune response, with sera containing antibodies specific for both Gc and whole CCHFV antigens (Fig. 3A).

**Figure 3.**
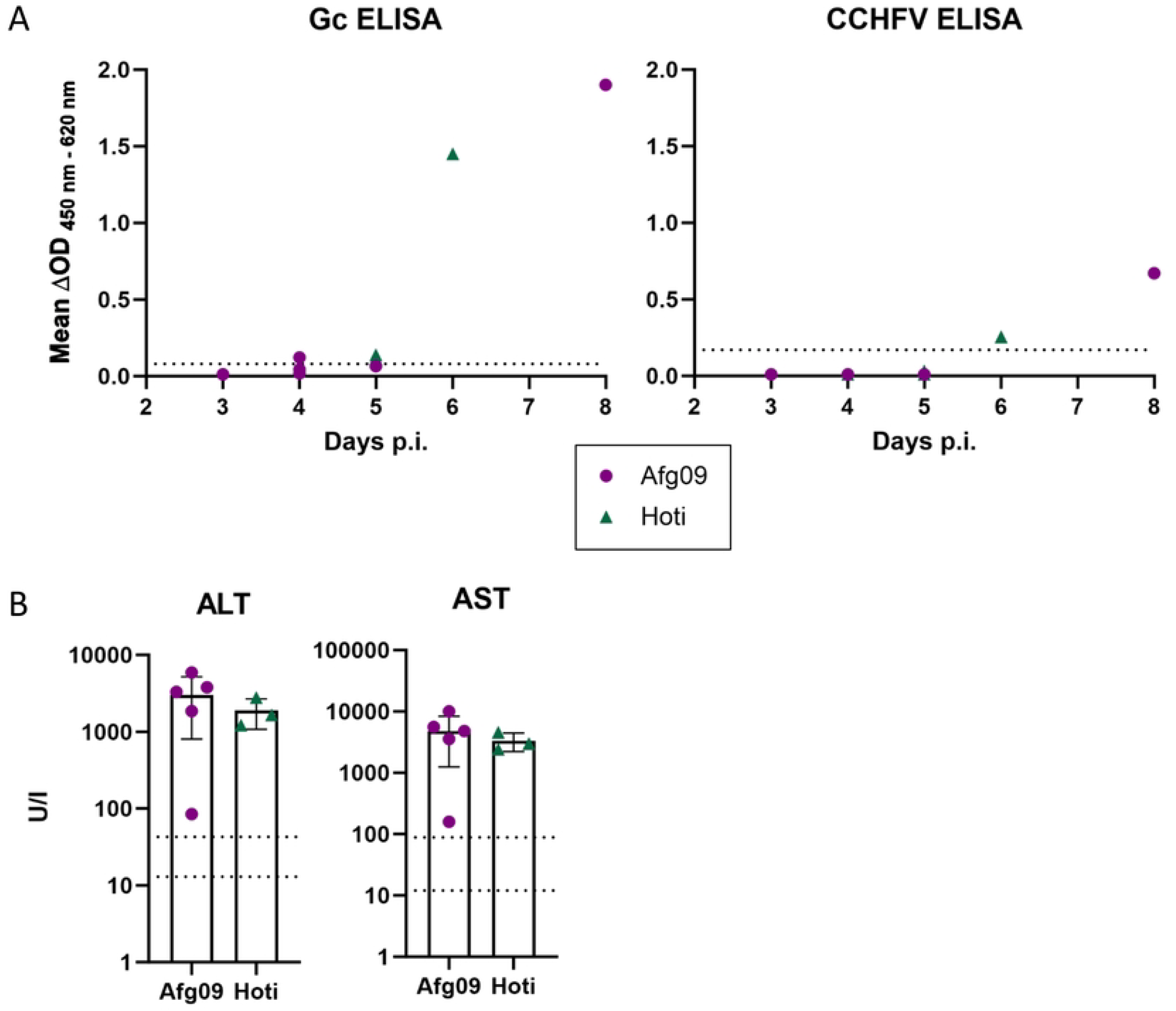
Serology of CCHFV Afg09 and Hoti-infected IFNAR^-/-^ mice. IFNAR^-/-^ mice were infected with either 100 TCID_50_ CCHFV Afg09 (purple) or Hoti (green). **A:** Final sera (Afg09 n = 6, Hoti n = 6) were analyzed with two virus-specific ELISAs using either inactivated CCHFV, measuring NP-specific antibodies or the major glycoprotein Gc for coating. Monoclonal antibodies detecting Gc or N protein were used as controls (not shown). Dotted lines indicate the respective lower limit of detection. **B:** Liver enzymes alanine aminotransferase (ALT) and aspartate aminotransferase (AST) were analyzed in final serum samples (Afg09 n = 5, Hoti n = 3). Dotted lines indicate the physiological range of healthy C57BL/6 mice(8).

Final serum samples were analyzed for the liver enzymes alanine aminotransferase (ALT) and aspartate aminotransferase (AST) as a sign of hepatocellular damage. Sera from five Afg09-infected and three Hoti-infected animals were available for analysis. This showed that ALT and AST values were well above the physiological reference range either after Afg09 or after Hoti infection (Fig. 3B).

Histopathological analysis of hematoxylin and eosin (H&E) stained slides revealed for both CCHFV isolates multifocal liver necrosis and lymphohistiocytic infiltrates (Fig. 4A, arrows). For the Afg09-infected mice, these necroses were generally more widespread and occasionally resulted in complete loss of the tissue architecture. Spleen histopathology indicated apoptosis in germinal follicle centers after CCHFV Afg09 infection and secondary follicles with only few apoptotic lymphocytes after CCHFV Hoti infection (Fig. 4A, arrowheads). Interestingly, nearly all organs examined, including the CNS with meninges, revealed *in situ* hybridization (ISH) staining without widespread and marked tissue damage (Fig. 4A). Quantification revealed that CCHFV genomes were detectable by ISH in 17% of the liver area in mice infected with CCHFV Afg09, compared to 3% in mice infected with CCHFV Hoti. In the spleen, viral genomes were detectable in approximately 9% of the tissue area in Afg09-infected mice, whereas only about 1% of the spleen area was CCHFV-positive in mice infected with Hoti. Similarly, in the CNS, CCHFV RNA was present in approximately 0.6% of the tissue area following Afg09 infection, compared to 0.1% in Hoti-infected animals (Fig. 4B). Altogether, ISH revealed a markedly higher abundance of viral RNA in tissues from mice infected with Afg09 compared to those infected with Hoti.

**Figure 4.**
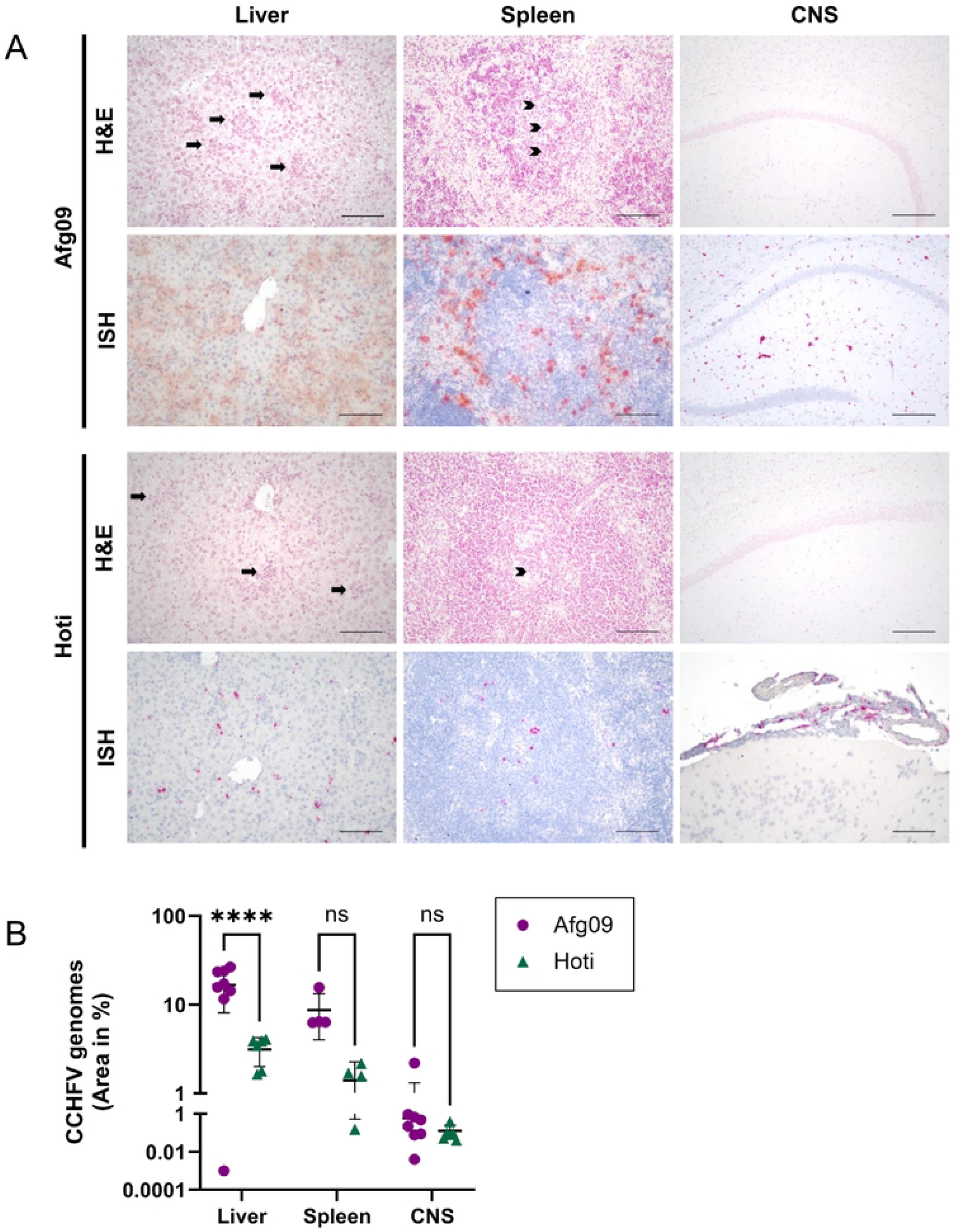
Pathology of CCHFV Afg09 and Hoti-infected IFNAR^-/-^ mice. Liver, spleen and CNS of CCHFV Afg09 (purple) or Hoti-infected (green) IFNAR^-/-^ mice were pathologically examined post-mortem. **A:** Shown are exemplary images of organ histopathology, H&E: hematoxylin and eosin staining. ISH: In situ Hybridization. Arrows indicate multifocal liver necrosis with loss of tissue architecture and lymphohistiocytic infiltrates. Arrowheads indicate apoptosis in germinal follicle centers. Scale bars: 100 µm. **B:** Quantification of ISH indicating percentage area of CCHFV-positive organ sections. Each data point represents a sample from an individual animal, horizonal lines represent the means ± SD. Datasets were analyzed using the Šídák’s multiple comparisons test. Asterisks indicate statistical significance as detailed between CCHFV Afg09 and Kosovo Hoti group: ∗∗∗∗p < 0.0001; ns = not significant.

## Discussion

In the present study, we established models for CCHFV Afg09 and Hoti based in C57BL/6J IFNAR^-/-^ mice. Infections with both viruses took a fulminant clinical course. In both mouse models, a clear, temporary increase in body temperature was measured 1-2 days before the clinical endpoint was reached. Fever is also one of the pre-haemorrhagic symptoms of CCHF in humans, as reviewed elsewhere(9). However, comparing the clinical course between the Afg09 and Hoti infection, clear differences were detected. Afg09-infected animals reached humane endpoint criteria between day 4-8 characterized by severe weight loss (>15%), reduced spontaneous behaviour and altered appearance as well as a preterminal drop in body temperature. Two of the eight animals succumbed during the course of the experiment. This is also reflected in the rapidly and dramatically worsening clinical score of 24.6 points at the time of euthanasia. These findings are consistent with other studies in which the Afg09 strain was lethal in IFNAR^-/-^ mice or when Interferon activity of wild type mice was counteracted by mouse-specific anti-IFNAR1 monoclonal antibodies(10–12). Hoti-infected mice reached the humane endpoint criteria on day 4-6 mainly because of a severe weight loss (>15%), with few other clinical signs. The final average clinical score was 11.2 points. In other studies using C57BL/6 IFNAR^-/-^ mice, Hoti-infected mice reached the endpoint 8 days after i.p. infection(13). The different criteria for the humane endpoint, >15% weight loss in our study and >25% in Hawman *et al*., 2018(13), may explain the difference in survival time. Notably, in a study by Hawman *et al*., 2019, subcutaneous infection with the CCHFV Hoti isolate even failed to result in a lethal outcome in the majority of infected IFNAR^−/−^ mice(14). Consistent with these findings, mice in which the type I interferon response was transiently blocked by using mouse-specific anti-IFNAR1 monoclonal antibodies also survived infection with the CCHFV Hoti isolate(11). This model has therefore been considered suitable for studying recovery following severe CCHFV disease. As humane endpoint criteria are defined individually by each laboratory in agreement with institutional animal welfare officers and local regulatory authorities, outcomes across animal models may vary accordingly. Nonetheless, a weight loss exceeding 15% is generally recognized as a severe clinical burden.

CCHFV infection in humans causes a systemic disease that affects multiple organs including the CNS and is frequently associated with a fatal outcome(9). In the present animal models, post-mortem examinations demonstrated that CCHFV genomes could be detected in all organs and sera examined, with markedly higher viral RNA levels in mice infected with Afg09 compared to those infected with Hoti. Infectious virus was detected in almost all organs tested and in the serum of Afg09-infected animals. Notably, the presence of virus in the CNS and eyes suggests a breach of the blood-brain and blood-ocular barriers, allowing viral entry into immune-privileged sites. In Hoti-infected mice, infectious virus was detected primarily in serum, thymus, lung and ovaries.

Previous studies had demonstrated infectious Afg09 in spleen, kidney, liver, heart, lung and CNS of infected mice(12). Our comprehensive analysis further identified viral presence in additional tissues, including thymus, eyes, mesenteric lymph nodes and the reproductive tract of both male and female mice. For Hoti, earlier investigations reported viral genomes in serum, liver, spleen, lymph nodes, lungs, CNS, eye and kidneys(13, 14). In the present study, we also detected viral RNA - and, in some cases, infectious virus – in the reproductive tract of both sexes. The relatively low levels of infectious virus in most organs of Hoti-infected mice supports the hypothesis that the mice are, in principle, capable of clearing the infection after a few days. This interpretation aligns with earlier findings in which mice infected with Hoti exhibited signs of recovery beginning on day 4 p.i.(14, 11). The detection of CCHFV-specific IgG antibodies in the sera from day 4 p.i. onwards further supports the activation of an adaptive immune response. Animals that remained in the study until day 6 or 8 p.i. exhibited both Gc-specific antibodies and also antibodies that react in the whole CCHFV ELISA. Concurrently, neither CCHFV genomes nor infectious virus were detectable in the sera of these animals. Furthermore, viral RNA levels and infectious viral titers in their organs were markedly lower compared to animals that had to be euthanized at earlier time points. Importantly, the detection of virus in the reproductive tract organs of female and male mice in both models suggests that CCHFV could also be transmitted sexually. This is consistent with individual human cases that provide evidence for sexual transmission in both male-to-female and female-to-male directions(15).

Organ damage resulting from infection with the CCHFV isolates Afg09 and Hoti was further characterized using blood chemistry and histopathological examinations. Blood chemistry analyses revealed significant elevations of liver enzymes (ALT and AST) in both mouse models compared to the reference range of uninfected C57BL/6 mice(8), indicating substantial liver damage following infection. Notably, enzyme levels were slightly higher in mice infected with Afg09 compared to those infected with Hoti. These findings align with previous studies suggesting comparable liver damage induced by both virus isolates(16).

Histopathological analysis confirmed liver damage consistent with serological findings, while spleen damage was less pronounced in Hoti-infected animals compared to those infected with Afg09. CNS abnormalities were observed only sporadically; however, viral RNA was detectable by ISH in all three analyzed organs. Generally, ISH staining in the CNS was weaker in Hoti-infected mice compared to Afg09-infected animals. The severity of tissue damage in individual organs tended to correlate positively with the detected viral load.

A previous study by Golden *et al*.(11) also investigated liver pathology in mice infected with these two CCHFV isolates, reporting comparable histopathological findings between isolates at day 4 p.i., consistent with our observations. In contrast to our in situ hybridization results, however, Golden and colleagues found similar viral RNA levels in the liver for both isolates at day 4. This discrepancy likely arises from differences in the timing of analyses; while Golden *et al*. euthanized animals at a fixed time point (day 4), animals in our study reached the clinical endpoint variably between days 4 and 6 p.i.. Additionally, Golden *et al*. demonstrated a significant decline in viral RNA by day 10 following Hoti infection, suggesting active viral clearance from day 4 onwards(11).

CCHFV poses a global health threat due to its widespread distribution and genetic diversity, which has resulted in the emergence of at least five distinct viral lineages. To ensure the development of broadly effective therapeutics and vaccines, it is critical to evaluate their efficacy across these diverse strains. To address this, we established side-by-side CCHFV mouse models using the Afg09 and Hoti isolates, representing the Asian and European lineages, respectively. Comparative analysis of these models revealed that both isolates cause lethal disease in IFNAR^-/-^ mice, although they differ in the severity and progression of illness.

## Methods

### Cells and viruses

Vero C1008 cells (ATCC CRL-1586) were cultured as described elsewhere(17). CCHFV Afghanistan-2990 (GenBank: HM452307.1, HM452306.1 and HM452305.1) and CCHFV Kosovo Hoti (GenBank: DQ133507, EU037902 and EU044832) were propagated and titrated in Vero C1008 cells. All experiments with CCHFV were carried out in the BSL4 laboratory of the Philipps University of Marburg, Germany.

### Animal experiments

Male and female C57BL/6J IFNAR^-/-^ mice (12-16 weeks old) carrying a knockout (KO) of the interferon alpha/beta receptor (IFNAR) were chosen as model because they are susceptible to infection with CCHFV(3). All experiments and protocols were approved by the local authorities (animal welfare committees; Marburg: Regierungspräsidium Gießen AZ V54 –19 c 20 15 h 01 MR 20/7 Nr. G 87/2022) and conducted according to the recommendations of Federation of European Laboratory Animal Science Associations (FELASA) and the Society for Laboratory Animal Science (GV-SOLAS) and were in compliance with the German animal welfare act as well as the directive 2010/63/EU. Tissue and blood samples from healthy, non-infected control mice was available from previous studies.

Two weeks prior to the start of the experiment, mice were assigned to the experimental groups (2-3 animals per sex) and housed in isocages (Tecniplast) under SPF conditions according to FELASA recommendations. One week before the experiment, they were tagged under brief isoflurane anesthesia (CP-Pharma) with an ear mark and implanted subcutaneously with a transponder (PTT-300, Plexx BV) for body temperature measurement. At the start of the experiment mice were infected intraperitoneally (i.p.) with 100 TCID_50_ CCHFV under brief isoflurane anesthesia. In addition to their normal diet, all animals were given access to a high-calorie energy gel after infection. The mice were monitored daily for weight, body temperature, general condition and spontaneous behaviour, with each category receiving a maximum score of 10. The sum of the four scores determined the clinical score. When a clinical score of ≥10, or ≥6 on two consecutive days, was reached, mice were euthanized by cervical dislocation under isoflurane anesthesia. 2 out of 8 Afg09-infected mice succumbed between days 3 and 4 despite daily monitoring and additional video surveillance before reaching the pre-defined human endpoint criteria. Consequently, an additional clinical examination was incorporated into subsequent studies on day 3 p.i. in the late afternoon. On days 3 and 7 p.i. or when the humane endpoint was reached, blood was drawn under brief isoflurane anesthesia by puncture of the facial vein.

### Quantitative real-time RT-PCR analysis of virus load in mouse tissue samples

Tissue and serum samples of mice were processed as described elsewhere(18). RT-qPCR was carried out using Altona diagnostics RealStar® CCHFV RT-PCR Kit 1.0 RUO (Cat. No. 181003) according to manufacturer’s instructions on a qTOWER^3^ (Analytik Jena).

### Virus titration by TCID_**50**_

To determine infectious virus in organ and serum samples, 10^4^ Vero C1008 cells were seeded in 96-well plates. The following day, cells were inoculated with 5-fold serial dilutions of supernatants from either infected cells or organ homogenates. At 6 days p.i. cytopathic effect (CPE) was analyzed via microscopy and TCID_50_/ml values were calculated according to Spearman and Kerber.

### Histopathological examination

Organs were collected after the humane endpoint was reached and processed for histological analysis as described elsewhere(19). To stain CCHFV RNA, either the Afg09-specific probe (Bio-Techne, RNAscope Probe - V-CCHFV-M-CPG-sense, Cat. No. 494801) or the Kosovo Hoti-specific probe (Bio-Techne, RNAscope Probe - V-CCHFV-M-env-sense, Cat. No. 497341) was used. Infected areas positive for CCHFV viral RNA by ISH were quantified by determining the relative area of tissue staining positive for viral RNA by automated image analysis of the whole section. Whole slides were scanned using the Hamamatsu NanoZoomer system. The images were then converted to 8-bit images using Image J and the threshold was adjusted to detect only positive areas. The measurements were set to determine ‘area’ and ‘area fraction’.

### CCHFV Gc-specific ELISA (IgG)

The CCHFV Gc ELISA was performed in the same way as our SARS-CoV-2 spike protein (S1) ELISA (IgG)(20). Deviations from the protocol were the following. Microtiter plates were coated with CCHFV glycoprotein (Gc) protein (The Native Antigen Company, REC31696) diluted to 0.5 µg/ml in phosphate buffered saline (PBS) and incubated for 20 hours at 4°C. For analysis, mouse sera were diluted 1:100 in PBS / 0.1% Tween®20 (PBST) with 1% powdered milk and allowed to react with the Gc protein for 1 hour. To confirm reliability and repeatability a blank, a negative control, and a Gc-specific mouse monoclonal antibody (11E7, dilution 1:50,000, BEI Resources, NIAID, NIH) were analyzed on each plate. Detection was performed using polyclonal goat anti-mouse immunoglobulins/HRP (Agilent DAKO; P044701-2, dilution 1:1,000, 30 min of incubation), 3,3’,5,5’-tetramethylbenzidine (TMB) substrate solution and TMB-Stop Solution. The optical density (OD) was determined at 450 nm – 620 nm using an automated spectrophotometer (PHOmo, Autobio Labtec Instruments Co., Ltd. or Synergy LX, Agilent BioTek). Each control and serum was analyzed in duplicate. The cut-off for a positive antibody response was calculated as the mean OD value of the results from 25 negative mouse sera plus four standard deviations.

### CCHFV whole virus ELISA (IgG)

To prepare CCHFV antigen for the ELISA, CCHFV strain Afg09-2990, (passage +3 upon receipt, GenBank accession numbers: HM452305.1, HM452306.1, HM452307.1) was used. Supernatants of CCHFV-infected Vero C1008 (ATCC CRL-1586) cells (MOI 0.01) were collected 3 days p.i. and clarified from cell debris. Viral particles were concentrated and purified by ultracentrifugation. Pellets were resuspended in PBS and inactivated by addition of a final concentration of 0.05% beta propiolactone (von 98.5%, pharma grade, Ferak Berlin GmbH) for 72 hours. The inactivation process has been verified. Mock antigen was prepared following the same procedure as for CCHFV antigen. Small volumes of antigen preparations were stored at -20°C until use. High binding single-break strip microtiter plates (Greiner bio-one, Cat.No.705074) were coated with 50 µl CCHFV or mock antigen (both diluted 1:50 in PBS) and incubated for 20 h at 4°C. Incubation times and buffers were used as described elsewhere for an Ebola virus antigen-based ELISA(21) Mouse sera and controls were diluted 1:100 in PBST containing 1% milk powder and allowed to react with CCHFV and mock antigen for 1 h. To confirm reliability and repeatability a blank, a negative control and a nucleoprotein-specific mouse monoclonal antibody (9D5, BEI Resources, NIAID, NIH) was used on each plate. Detection was performed as described above for the Gc-specific ELISA. Each control and serum was analyzed once and the OD value of each sample on mock antigen was substracted from the OD value on CCHFV antigen to obtain corrected OD values (ΔOD). The cut-off for a positive antibody response was calculated as the mean OD value of the results from 25 negative mouse sera plus four standard deviations.

### Clinical serum chemistry

Serum clinical chemistry was assessed using the Piccolo Xpress Chemistry Analyzer and the General Chemistry 13 panel (both Abaxis, Union City, CA, USA) to determine levels of the liver enzymes aspartate aminotransferase (AST) and alanine aminotransferase (ALT) as an indicator of hepatocellular injury according to the manufacturer’s instructions.

### Statistical analysis

The statistical analysis was carried out using Graphpad Prism 10 software.

## Data availability

Data sets are available upon reasonable request.

## Acknowledgements

We thank all individuals who contributed to this work. We are particularly grateful to U. Kalinke (TiHo, Hannover, Germany) for kindly providing C57BL/6J IFNAR^−/−^ mice. We also thank the Institute of Veterinary Pathology (JLU, Giessen, Germany) for providing technical equipment and for the preparation of histological sections. Special thanks go to the BSL-4 team, especially G. Ludwig and S. Schmidt, whose support was essential for the successful completion of this work. This publication was supported by kind provision of the CCHFV Kosovo Hoti isolate by the European Virus Archive Global (EVA-GLOBAL) project which has received funding from the European Union’s Horizon 2020 research and innovation programme under grant agreement No 871029. The following reagents were obtained from the Joel M. Dalrymple – Clarence J. Peters USAMRIID Antibody Collection through BEI Resources, NIAID, NIH: Monoclonal Anti-Crimean-Congo Hemorrhagic Fever Virus Pre-Gc Glycoprotein, Clone 11E7 (produced in vitro), NR-40277 and monoclonal Anti-Crimean-Congo Hemorrhagic Fever Virus Nucleocapsid Protein, Clone 9D5 (produced in vitro), NR-40270. The project was funded by DZIF TTU Emerging Infections (FKZ 8033801809).

## Author contributions

Conceptualization, C.R., V.K., A.K.; methodology, C.R., V.K., A.K.; formal analysis, C.R., V.K., A.K.; investigation, all authors; writing—original draft preparation, C.R., V.K., A.K.; writing— review and editing, all authors.; visualization, C.R., V.K., A.K.; supervision, S.B., V.K., A.K.; funding acquisition, S.B., V.K., A.K.

